# Structural basis for ultrapotent neutralization of human metapneumovirus

**DOI:** 10.1101/2022.03.14.484292

**Authors:** Avik Banerjee, Jiachen Huang, Scott A. Rush, Jackelyn Murray, Aaron D. Gingerich, Fredejah Royer, Ching-Lin Hsieh, Ralph A. Tripp, Jason S. McLellan, Jarrod J. Mousa

**Affiliations:** Center for Vaccines and Immunology, College of Veterinary Medicine, University of Georgia, Athens, GA 30602; Department of Infectious Diseases, College of Veterinary Medicine, University of Georgia, Athens, GA 30602; Department of Molecular Biosciences, The University of Texas at Austin, Austin, TX 78712; Department of Biochemistry and Molecular Biology, Franklin College of Arts and Sciences, University of Georgia, Athens, GA 30602

## Abstract

Human metapneumovirus (hMPV) is a leading cause of morbidity and hospitalization among children worldwide, however, no vaccines or therapeutics are currently available for hMPV disease prevention and treatment. The hMPV fusion (F) protein is the sole target of neutralizing antibodies. To map the immunodominant epitopes on the hMPV F protein, we isolated a panel of human monoclonal antibodies (mAbs), and the mAbs were assessed for binding avidity, neutralization potency, and epitope specificity. We found the majority of the mAbs target diverse epitopes on the hMPV F protein, and we discovered multiple mAb binding approaches for antigenic site III. The most potent mAb, MPV467, which had picomolar potency, was examined in prophylactic and therapeutic mouse challenge studies, and MPV467 limited virus replication in mouse lungs when administered 24 hrs before or 72 hrs after viral infection. We determined the structure of MPV467 in complex with the hMPV F protein using cryo-electron microscopy to a resolution of 3.3 Å, which revealed a complex novel prefusion-specific epitope overlapping antigenic sites II and V on a single protomer. Overall, our data reveal new insights into the immunodominant antigenic epitopes on the hMPV F protein, identify a new mAb therapy for hMPV F disease prevention and treatment, and provide the discovery of a unique pre-fusion-specific epitope on the hMPV F protein.

## Introduction

Human metapneumovirus (hMPV) is a leading cause of respiratory disease in children and the elderly.^1–5^ Initially identified in 2001 in samples collected from children with respiratory tract infection in the Netherlands^6^ the clinical features of hMPV are similar to those of respiratory syncytial virus (RSV), and include mid-to-upper respiratory tract infection that may require hospitalization.^7^ Severe disease can occur in immunocompromised patients, such as those undergoing lung transplant^8^ hematopoietic stem cell transplant^9–12^ as well as those living with HIV^13^ and COPD.^14^ There are no approved vaccines or specific treatments available for hMPV infection, in contrast to RSV, for which palivizumab^15^ has been in use for many years in specific high-risk infant groups.

Serological studies have shown that nearly all children are seropositive for hMPV by five years of age.^16^ hMPV has three surface glycoproteins – the small hydrophobic (SH), attachment (G), and fusion (F) proteins. Of these, the hMPV F protein is the only target of neutralizing antibodies^17^ which is different than RSV where both the RSV G and RSV F proteins elicit neutralizing antibodies^18^. There are no licensed vaccines to protect against hMPV, but several candidates have been examined in animal models, including live-attenuated viruses, recombinant viruses, vectored vaccines, and recombinant surface proteins.^19^ Limited vaccine candidates have advanced to clinical trials, including a live-attenuated hMPV vaccine (NCT01255410), and more recently, an mRNA-based vaccine combined with parainfluenza virus 3 (NCT03392389, NCT04144348). Similar to vaccine enhanced disease observed with formalin-inactivated RSV^20–22^ vaccines using formalin-inactivated hMPV result in enhanced disease following viral infection in mice, cotton rats, and macaques.^23,24^

The hMPV F protein is a trimeric class I viral fusion protein that has high conservation between viral subgroups (A1, A2, B1, B2).^25^ hMPV can infect respiratory epithelial cells in the absence of the hMPV G protein, although hMPV G is required for viral fitness *in vivo*.^26^ The hMPV F protein contains an RGD motif, and the receptor has been hypothesized to be α_5_β_1_ integrin.^27^ Heparan sulfate has also been shown to have a role in hMPV F protein-mediated attachment^28^ and we recently demonstrated direct binding between heparan sulfate and the hMPV F protein.^29^ hMPV F induces fusion of viral and host cell membranes in a transition from the metastable pre-fusion state to the post-fusion conformation.^30^ X-ray crystal structures of the hMPV F protein in the pre-fusion^31^ and post-fusion^29,32^ conformations have been elucidated, and the protein shares similar structural topology with the RSV F protein.^33^

There has been a paucity of information regarding specific epitopes on the hMPV F protein compared to RSV F. Clear differences in the immunologic features between RSV F and hMPV F have been identified. For example, pre-fusion and post-fusion hMPV F proteins elicit similar antibody responses, suggesting that the majority of the neutralizing epitopes are present in both conformations^29,31,34^ while for RSV F the majority and the most potent neutralizing antibodies target pre-fusion-specific epitopes.^25,35,36^ Known neutralizing epitopes on the hMPV F protein include antigenic sites IV^32,37,38^ III^39,40^ and V^41^ based upon identification of RSV F mAbs that also neutralize hMPV F. A unique epitope targeted by the human mAb DS7 has been structurally defined.^42^ Additionally, we recently discovered a novel epitope located within the trimeric interface of the hMPV F protein, which was defined by mAb MPV458.^43^ To further our understanding of neutralizing hMPV F epitopes, we isolated a panel of 18 new human mAbs to the hMPV F protein. We discovered two mAbs in particular, MPV467 and MPV487, have potent neutralizing activity, with MPV467 being exceptionally potent. We determined that MPV467 can prevent and treat hMPV infection in mice, and we determined using cryo-electron microscopy that the epitope of MPV467 targets a complex binding site interfacing both antigenic sites II and V.

## Results

### Human monoclonal antibody sequence determinants

To further define the antigenic epitopes on the hMPV F protein, we isolated 18 new human mAbs using our previously described hMPV B2 F protein^29^ with sixteen mAbs generated via human hybridoma technology while two were derived from antigen-specific single B cell sorting (MPV491, 503). The antibody-encoding genes were sequenced, and the results indicated the usage of a diverse set of immunoglobulin *V* genes across the entire panel **(Figure 1A, Table S1)**. mAbs utilizing the VH1-69 gene were the most abundant. VH3 and VH4 gene families comprised the majority of the additional mAbs. Diversity was also present in the light chain, with eight and five unique genes utilized for the kappa and lambda mAbs, respectively. Kappa isotype mAbs utilized VK1, VK2, VK3, and VK4 gene families, while lambda isotype mAbs used VL1 and VL3 gene families. The lengths of the heavy and light chain junctions ranged from 14 to 23 amino acids for the heavy chain, 8 to 10 amino acids for the kappa chain, and 9 to 11 amino acids for the lambda chain **(Figure 1B)**. The percent identities of the variable genes to the germline sequence ranged 88–97% for the heavy chain (93.6% average) and 90–97% for the light chain (93.7% average) **(Figure 1C)**.

**Figure 1.**
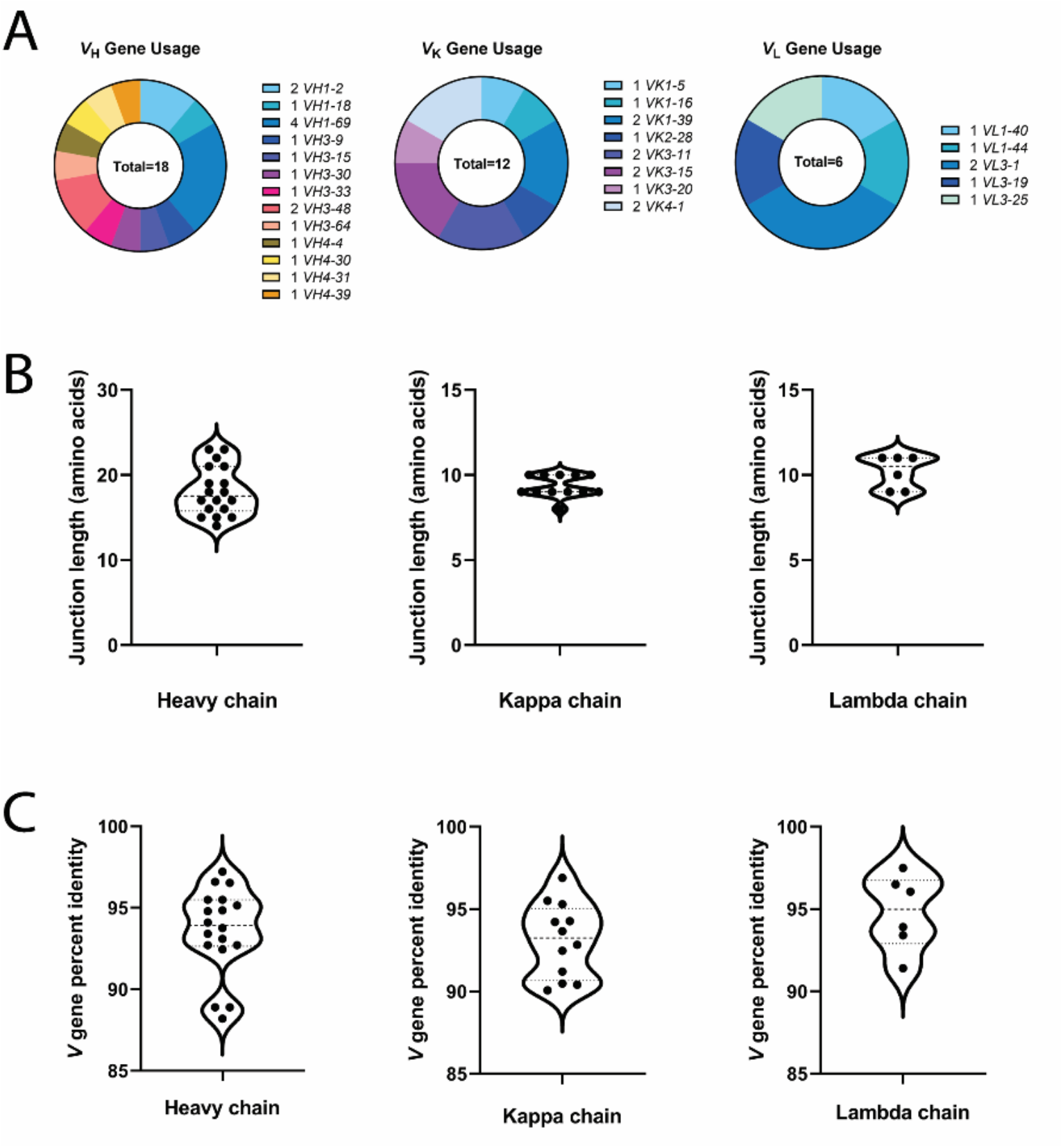
Sequence determinants of the isolated mAbs. (A) The usage of heavy, kappa, and lambda chain genes are shown as a proportion of all respective genes from the panel of isolated mAbs. (B) The amino acid lengths of the junction for the heavy and light chains are shown. (C) The percent identity of the *V* gene to predicted germline sequences are shown.

### mAb binding and functional properties

The neutralizing activity of each mAb was determined by plaque-reduction assay using representative viruses from each genotype of hMPV, i.e., hMPV CAN/97-83 (genotype A) and hMPV TN/93-32 (genotype B) **(Table 1, Figure S1)**. All mAbs had neutralizing activity against viruses from both genotypes, with mAbs MPV467, MPV487, MPV454, MPV482, and MPV488 having neutralizing activity below 20 ng/mL against hMPV CAN/97-83. mAbs MPV467, MPV487, and MPV454 have the most potent neutralizing activity against both hMPV CAN/97-83 and hMPV TN/93-32, with MPV467 reaching picomolar activity (below 1 ng/mL) against hMPV TN/93-32. mAbs MPV86, MPV488, MPV485, and MPV477 demonstrated preferential neutralization of hMPV CAN/97-83 based on at least a 20-fold lower IC_50_ compared to hMPV TN/93-32. The binding properties of the mAbs were assessed using a panel of hMPV F proteins from each subgroup (hMPV A1 F, hMPV A2 F, hMPV B1 F, hMPV B2 F) containing mixtures of pre-fusion and post-fusion hMPV F. mAb binding to additional constructs containing exclusively monomeric pre-fusion hMPV F, post-fusion hMPV F, and a predominantly trimeric pre-fusion hMPV F (hMPV B2 F GCN4) was also assessed. Several binding patterns were observed. MPV487, MPV482, MPV503, MPV414, MPV86, and MPV488 had limited binding to post-fusion F constructs and favored binding to pre-fusion constructs. mAbs MPV467, MPV454, MPV477, MPV486, and MPV464 bound to both pre-fusion and post-fusion constructs, but had higher binding affinity for pre-fusion proteins. mAbs MPV478, MPV483, MPV481, MPV456, MPV491, MPV489, and MPV485 bound equally to both pre-fusion and post-fusion constructs.

**Table 1.** hMPV F-specific mAb neutralization and binding properties.

| Table 1. hMPV F-specific mAb neutralization and binding properties. |  |  |  |  |  |  |  |  |  |  |  |  |
| --- | --- | --- | --- | --- | --- | --- | --- | --- | --- | --- | --- | --- |
| mAb | IC<sub>50</sub> (ng/mL) |  | EC<sub>50</sub> (ng/mL) |  |  |  |  |  |  |  |  |  |
|  | hMPV CAN/97-83 | hMPV TN/93-32 | hMPV A1 F | hMPV A2 F | hMPV B1 F | hMPV B2 F | hMPV A1 F monomer | hMPV A1 F trimer (heated) | hMPV B2 F monomer | hMPV B2 F trimer (heated) | hMPV B2 F GCN4 | RSV A2 F DsCav1 |
| 467 | 5 | 0.6 | 380 | 120 | 120 | 370 | 53 | 260 | 60 | 650 | 61 | > |
| 487 | 3 | 12 | 110 | 100 | 65 | 370 | 130 | > | 99 | 540 | 71 | > |
| 454 | 2 | 25 | 490 | 120 | 120 | 5200 | 52 | 140 | 46 | 150 | 68 | > |
| 482 | 14 | 47 | 130 | 57 | 72 | 330 | 290 | 350 | 200 | > | 92 | > |
| 486 | 510 | 62 | 53 | 43 | 100 | 230 | 88 | 220 | 110 | 240 | 100 | > |
| 478 | 210 | 68 | 59 | 21 | 22 | 27 | 12 | 24 | 20 | 20 | 20 | > |
| 483 | 32 | 110 | 120 | 70 | 120 | 220 | 160 | 270 | 130 | 330 | 140 | > |
| 503 | 330 | 190 | 3600 | > | > | 3400 | 1200 | > | 71 | > | 61 | > |
| 481 | 96 | 260 | 310 | 110 | > | 160 | 170 | 280 | 200 | 310 | 110 | > |
| 464 | 140 | 400 | 280 | 83 | 86 | 980 | 67 | 240 | 98 | 290 | 120 | > |
| 414 | 44 | 590 | 150 | 34 | 34 | 1800 | 74 | > | 190 | > | 61 | > |
| 456 | 450 | 810 | 1400 | 420 | 490 | 610 | 3500 | 1800 | > | 1800 | 690 | > |
| 86 | 57 | 1800 | 300 | 83 | 73 | 5700 | 180 | > | 210 | > | 120 | > |
| 491 | 5700 | 1900 | 1500 | 2200 | 2000 | 2100 | 1300 | 1100 | 480 | 1700 | 500 | > |
| 489 | 3600 | 2300 | 5000 | 9300 | > | 4200 | 7000 | 4500 | 1300 | 18000 | 2300 | > |
| 488 | 18 | 2400 | > | > | > | > | > | > | 63 | > | 73 | > |
| 485 | 91 | 2600 | 110 | 86 | 78 | 180 | 240 | 320 | 180 | 360 | 160 | > |
| 477 | 54 | 2900 | 83 | 23 | 20 | 63 | 49 | 87 | 210 | 63 | 37 | > |
| Neutralization values were determined using a plaque-reduction assay, where the IC<sub>50</sub> corresponds to the mAb concentration at which 50% plaque reduction was observed. EC<sub>50</sub> values correspond to the concentration at which the half-maximum signal was obtained in ELISA, based on the optical density at 405 nm. > indicates the binding signal was below 1 µg/mL at the highest mAb concentration tested. Each value is an average of three technical replicates for neutralization experiments and four technical replicates for binding experiments. Each experiment was repeated independently at least twice. |  |  |  |  |  |  |  |  |  |  |  |  |

### Epitope mapping

To determine the general binding epitopes for the panel of 18 mAbs, we conducted an epitope binning experiment using biolayer interferometry as previously described.^43,44^ Biosensors were loaded with hMPV B2 pre-fusion F protein, associated with a test mAb, and then exposed to a control mAb with a known epitope to determine competition profiles **(Figure 2A)**. Control mAbs targeting known hMPV epitopes included mAbs MPE8 and MPV364 (site III), DS7 and MPV196 (DS7 epitope), 101F (site IV), and MPV458 (66-87 intratrimer epitope) **(Figure 2B)**. MPV481 and MPV483 were mapped to antigenic site IV, while MPV454 competed with both 101F and DS7, suggesting it binds an intermediate epitope between site IV and the DS7 site **(Figure 2B)**. However, MPV454 did not compete with the previously discovered MPV196, which competes with DS7. MPV464, MPV491, MPV485, and MPV477 competed with both MPE8 and DS7 similar to our previous results with MPV196, MPV201, and MPV314.^44^ mAbs MPV86, MPV414, MPV482, MPV487, and MPV503 were “MPV364-like”, competing with MPE8 and MPV364 but not with DS7. This differential binding mode at antigenic site III has been previously defined by competition or the lack thereof with DS7.^44^ No mAbs were observed to compete with the intratrimer-targeting MPV458, although intermediate competition was observed between several mAbs and MPV458, suggesting that these may partially block binding of MPV458 or limit exposure of the intratrimer epitope bound by MPV458 centered at amino acids 66-87. MPV488 and MPV489 had partial competition with MPE8 but not MPV364, while MPV467 and MPV456 had partial competition with MPV364 but not MPE8, suggesting additional epitopes are present near antigenic site III. MPV486 and MPV478 showed partial competition with nearly all mAbs and their epitope could not be defined.

**Figure 2.**
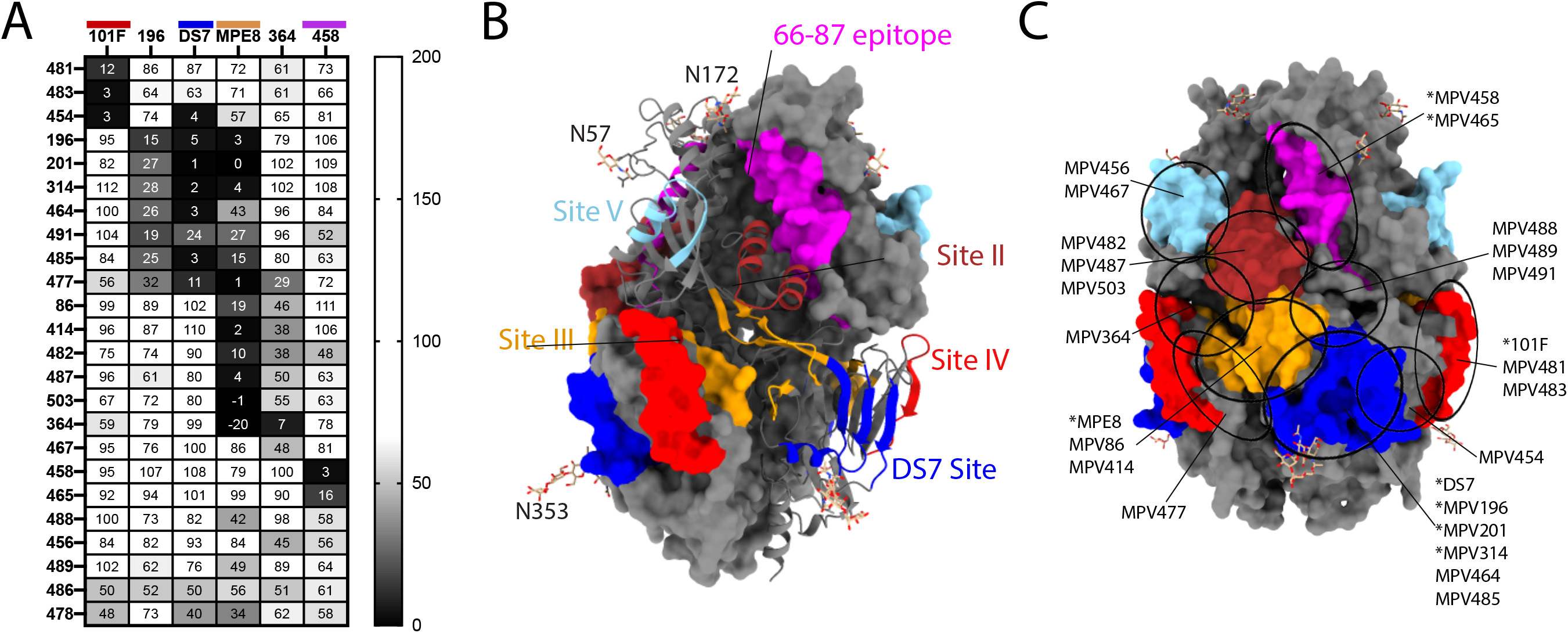
Epitope mapping of the hMPV F-specific mAbs. (A) Epitope binning for mAbs binding to the hMPV B2 F protein. Data indicate the percent binding of the second antibody in the presence of the primary antibody, as compared to the second antibody alone. Cells are colored in a gradient according to the legend displayed right. (B) Epitopes for control mAbs 101F (site IV), 196 and DS7 (DS7 epitope), MPE8 and MPV364 (site III), and MPV458 (66-87 epitope) were used as the second mAb and are labeled according to the colors in (A). Select control mAbs were also used as the first mAb as positive blocking controls. (C) The mAb binding sites of MPE8 and MPV364 (site III), 101F (site IV), DS7, and MPV458 (66-87 epitope) are displayed on the surface of monomeric prefusion hMPV F. Estimated epitopes from epitope binning for each mAb are displayed. mAbs with an asterisk were previously discovered.

### hMPV F-specific mAbs enhance antibody-dependent phagocytosis of THP-1 cells

Antibodies that bind to different hMPV F antigenic sites were selected to evaluate antibody-dependent phagocytic activities **(Figure 3)**. All the mAbs tested significantly enhanced the phagocytosis of THP-1 cells compared to the blank and isotype control mAb (PhtD3, a mAb that binds to *Streptococcus pneumoniae*) controls *in vitro*. Antibodies that bind to the DS7-site showed overall higher phagocytic scores while the rest of the mAbs varied in ADP activity. In addition, no correlation was observed between the EC_50_/IC_50_ and the ADP activity of the mAbs, suggesting the ADP activities of hMPV F-specific mAbs are independent from the Fab binding epitopes. These data suggest additional protective mechanisms of anti-hMPV F mAbs beyond neutralization.

**Figure 3.**
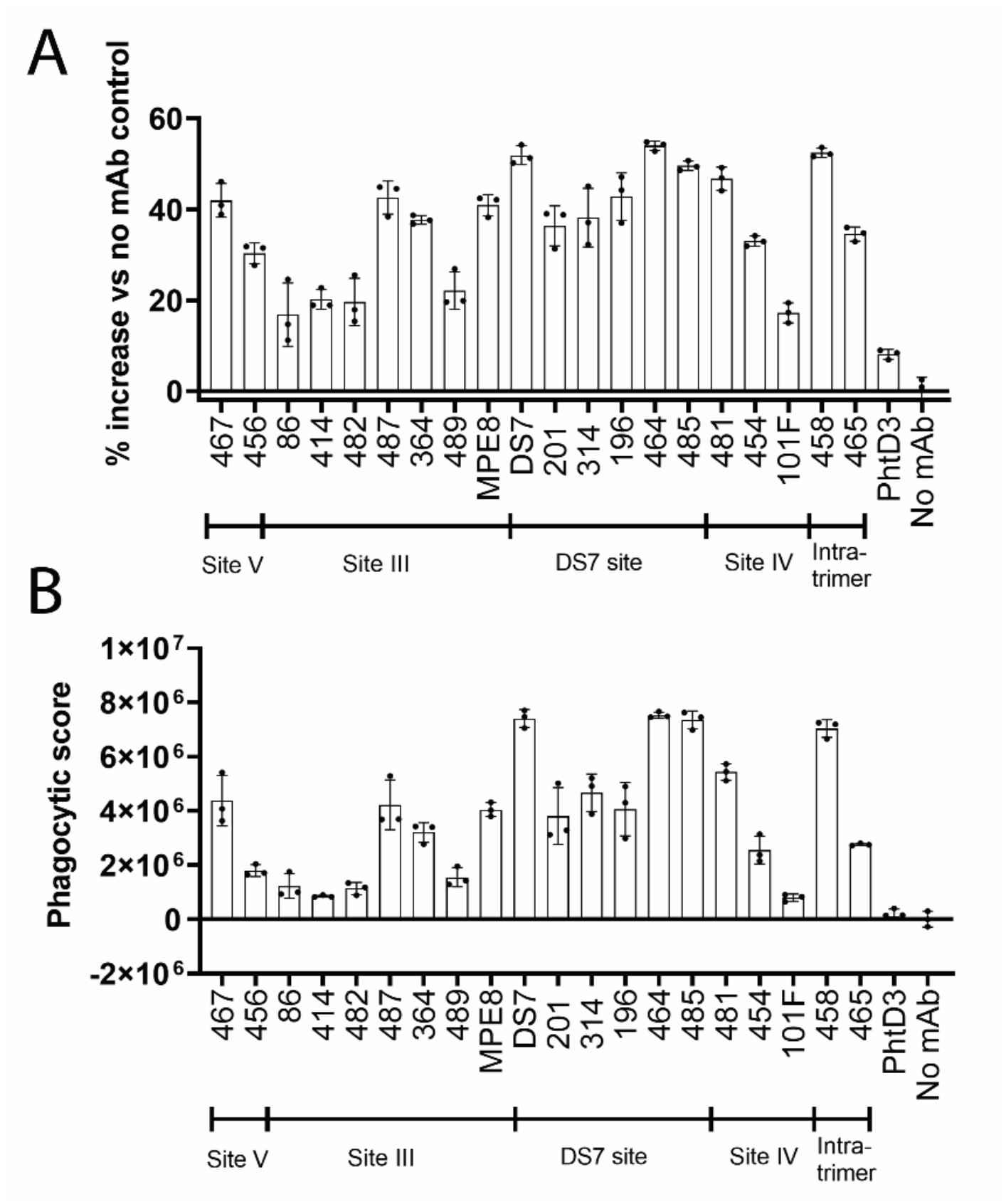
The percent phagocytosis of hMPV F-coated beads by THP-1 cells in the presence of each mAb was assessed using flow cytometry. The relative percent increase of phagocytic cells for each mAb relative to the no mAb control (A), in addition to the phagocytic score (B), are shown. Bars represent the average of three replicates, while errors bars are the standard deviation.

### Therapeutic efficacy of MPV467

MPV467 was the most potent mAb of the panel, reaching picomolar neutralization potency against hMPV TN/93-32 and potently neutralizing hMPV CAN/97-83. This mAb has pre-fusion preference properties, as substantial binding is lost to post-fusion hMPV F proteins **(Table 1)**. Based on these data, we tested the protective efficacy of mAb MPV467 in an hMPV infection model in BALB/c mice. Male and female mice were treated with PBS, an isotype control human mAb, or mAb MPV467 24 hrs prior to hMPV infection or three days after hMPV infection in both prophylactic and treatment studies **(Figure 4)**. On day five, viral titers in the lungs of mice were determined by plaque assay. No virus was detectable in mice treated with MPV467 in either study, while virus was present in both PBS-treated and isotype-mAb-treated mice. No difference was observed between MPV467 and uninfected mice, and no difference was observed between PBS and isotype control mice in either study.

**Figure 4.**
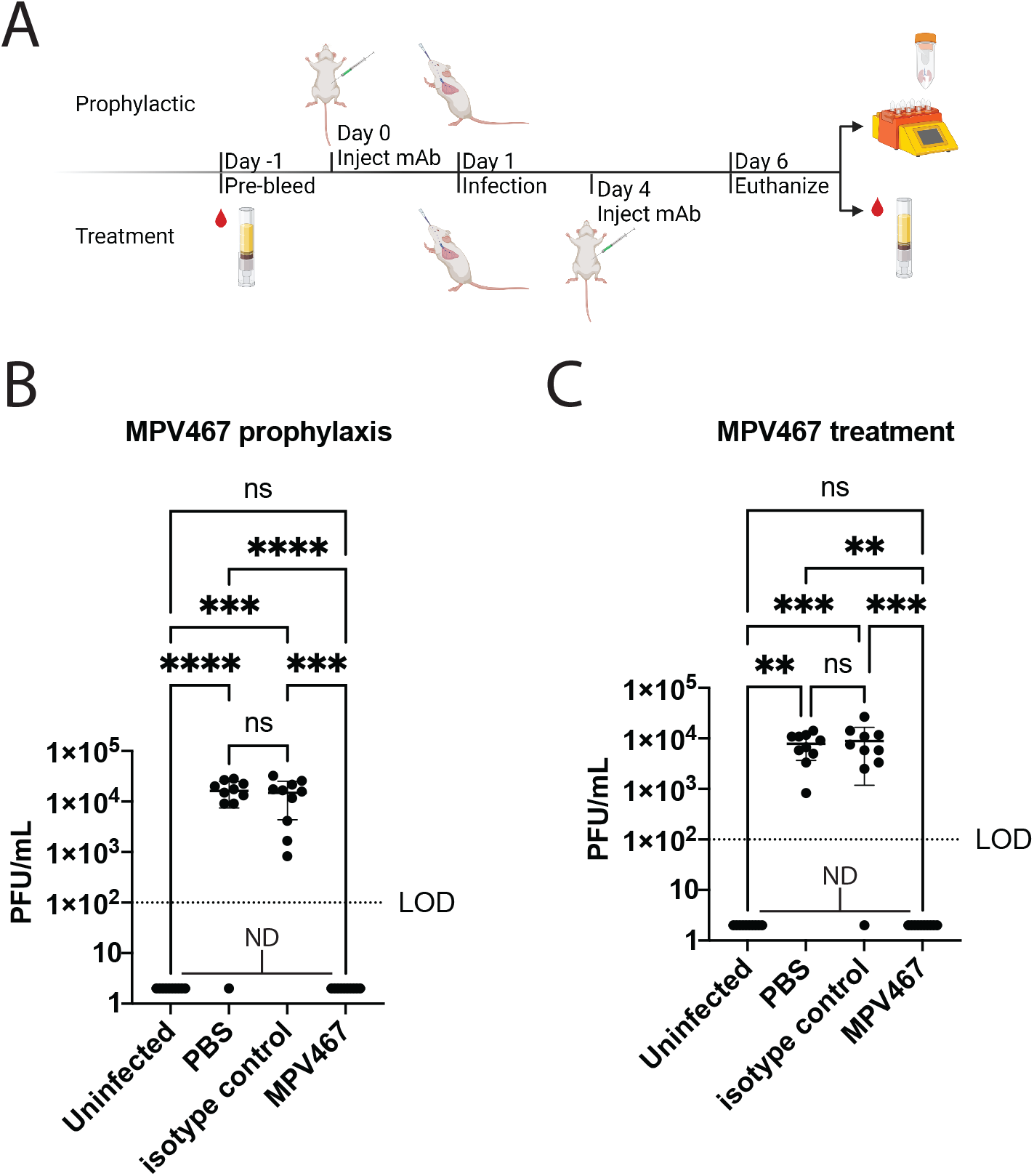
Protective efficacy of MPV467 against hMPV replication in *vivo*. (A) BALB/c mice were treated intraperitoneally with 10 mg/kg of mAb MPV467 24 hrs prior (prophylaxis study) or three days after (treatment study) intranasal hMPV infection. Viral titers in the lung homogenates of BALB/c mice in each treatment group (n = 10 mice per group, 5 males, 5 females) in prophylaxis study (B) and treatment study (C) were determined by plaque assay. n.s. = not significant, **P = 0.0016, ***P = 0.0003-0.0001, ****P <0.0001. Limit of detection (LOD) is indicated with a dashed line. ND=not detected.

### Structural definition of the hMPV F-MPV467 complex

As MPV467 is the most potent hMPV F mAb described to date, is protective against and can treat hMPV infection, and targets an undefined epitope, we determined the structure of MPV467 in complex with pre-fusion hMPV F using cryo-electron microscopy (cryo-EM) to a global resolution of 3.3 Å **(Figure 5)**. A final map for model building was generated by particle subtraction of the flexible Fab constant regions and sharpening via DeepEMhancer.^45^ The structure reveals that MPV467 has an angle of approach directed down toward the viral membrane **(Figure 5A, 5B)**. The heavy and light chains bury a surface area of 567 Å^2^ and 322 Å^2^, respectively, on hMPV F. The MPV467 epitope spans antigenic sites II and V, but also contacts a single residue in antigenic site III, Tyr44. The hMPV F site V residue Arg156 makes numerous interactions to MPV467 via hydrogen bonds to both CDRH1 and CDRH3 mainchain atoms as well as a salt bridge interaction with CDRH1 Asp31 **(Figure 5C)**. Additionally, CDRH1 Asp31 also forms a hydrogen bond with site V residue Thr150. The conformationally immobile antigenic site II is bound by the MPV467 CDRH3 through two hydrogen bond interactions with hMPV F Asn233 and Thr236. The only specific interaction between the light chain of MPV467 and hMPV F is through a mainchain hydrogen bond between CDRL1 Asn30 and hMPV F Ala238.

**Figure 5.**
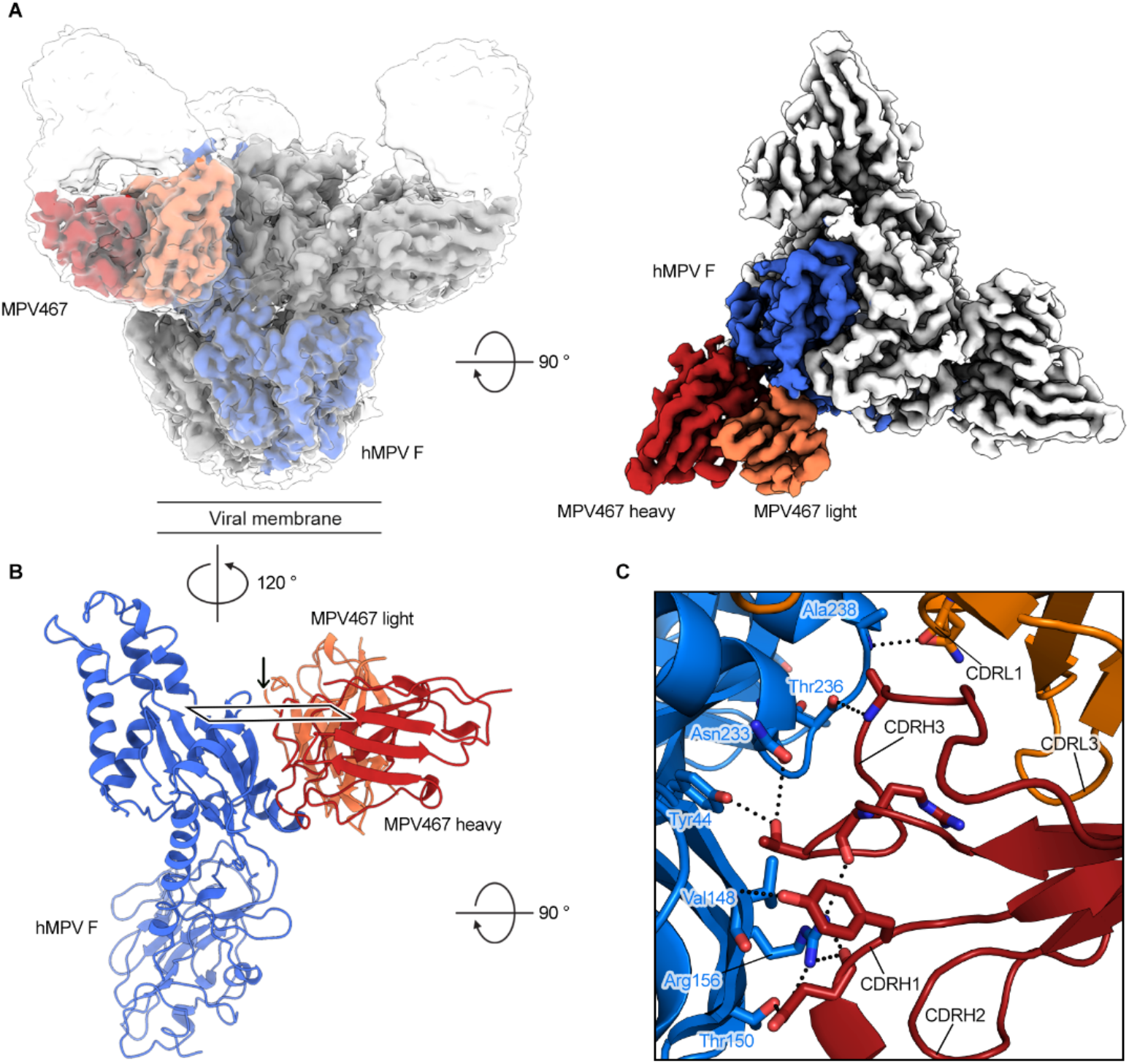
MPV467 binds pre-fusion hMPV F at sites II and V. (A) (left) Side view of the hMPV F–MPV467 Fab complex cryo-EM map shown at two different contour levels. Global map shown as white transparent map. Particle-subtracted, DeepEMhanced map is opaque with a single protomer colored (hMPV F: blue, Fab: red/orange). (right) Top-down view of particle-subtracted, DeepEMhanced map. (B) A single protomer of the hMPV F trimer and MPV467 Fab variable domain are shown as ribbons (hMPV F: blue, Fab: red/orange). (C) Zoomed in view of the MPV467 interface with hMPV F. View direction as shown by box and arrow in panel (B). Important residues shown as sticks. Hydrogen bonds and salt bridges depicted as black dotted lines. Oxygen atoms are colored red and nitrogens are blue.

## Discussion

Humans have been exposed to hMPV for at least 70 years^6^ yet the predominant epitopes on the hMPV F protein have remained elusive. We and others have previously isolated human mAbs to the hMPV F protein and have begun to map the dominant antigenic epitopes targeted by neutralizing mAbs. We discovered the repertoire, binding, and neutralizing capacity of 18 new human mAbs, and these mAbs targeted multiple epitopes, including sites IV, DS7, and at least four unique epitopes that bind at or near antigenic site III.

Variable region sequence analysis showed the diversity of V_H_ and V_K_/V_L_ gene usage in hMPV F-specific mAbs. Among these genes, V_H_1-69 is shared by 4 out of 18 mAbs (MPV86, MPV414, MPV483, and MPV503), and three of them (except for MPV483) showed similar binning profiles that compete with MPE8 and MPV364 **(Figure 2A, 2B)**, indicating they bind to the same hMPV F epitope possibly via similar binding patterns of the heavy chains. Similar correlations between binding epitopes and V gene usage were observed in light chains as well. Both mAbs MPV86 and 487 that share IGKV3-11 and mAbs MPV 414 and 503 that share IGKV3-15 bind to the MPV364 site, while mAbs MPV464 and 485 that share IGLV3-1 bind to DS7 site. mAbs MPV414 and MPV503 were identified from two different subjects using two different approaches, yet they have the same set of V_H_/V_K_ genes, suggesting this pair of genes might be favored by hMPV F MPV364 site–specific mAbs. This finding will need to be further investigated with a larger pool of mAbs in the future.

With our previously reported mAbs^43,46^ and the new batch of hMPV F-specific mAbs in this study, we are able to extensively map the major antigenic sites on hMPV F. In addition to the known antigenic sites III, IV, the DS7-site, and the 66-87-site, we further characterized site V and identified a putative site II. Like site III and site IV, the positions of both site V and site II resemble their counterparts on RSV F, suggesting the epitopes in these areas share structural features that can be recognized by human antibodies. However, no mAbs were found to bind the counterparts of RSV F site Ø on hMPV F, further suggesting mAbs to this epitope may be limited due to N-linked glycans on hMPV F at site Ø (Asn57 & Asn172) as previously suggested.^31^

The most potent mAb, MPV467, binds an epitope located across antigenic sites II and V. Structures of hMPV F in its pre- and post-fusion conformations reveal that it has structural homology with the related RSV F protein and neutralizing epitopes on RSV F can be expected to have counterparts on hMPV F. Structurally, antigenic sites Ø and V are pre-fusion specific and previously isolated RSV antibodies and structural studies have shown that antibodies targeting these regions tend to be highly potent neutralizers.^35,47^ Here with the cryo-EM structure of mAb MPV467 in complex with hMPV F, we determined this mAb is one of the first site V-targeting antibodies discovered for hMPV F and that it does indeed share the characteristic of potent neutralization. MPV467 binds the beta hairpin (β3-β4) located within antigenic site V, which undergoes a large conformational change during the transition from pre-fusion to post-fusion. This region of the MPV467 epitope is likely responsible for the potency of MPV467. While bound, the fusion protein cannot transition to the post-fusion conformation, which is essential for efficient viral infection. Additionally, the helix-turn-helix (α6-α7) located within antigenic site II that is bound by MPV467 does not undergo a conformational change between pre- and post-fusion states. The binding of residues in this region is likely why we see the ability of MPV467 to bind both conformational states. Since only this part of the epitope is present in the post-fusion conformation, we see a drop in binding affinity compared to the pre-fusion conformation which includes the entire epitope.

An RSV/hMPV F cross-reactive mAb M1C7^41^ was previously reported as a potent neutralizing mAb targeting site V. Similarly, an RSV F site V-specific mAb, hRSV90^47^ also has a low IC_50_ against RSV (4 ng/mL for RSV A, 10 ng/mL for RSV B) indicating site V is a vulnerable area that is favored by ultrapotent neutralizing mAbs for Pneumoviruses. The site V epitope of both RSV F and hMPV F is located on the N-terminus of the F_1_ subunit, closely following the fusion peptide, which is buried in the center of the trimeric pre-fusion F protein. Hence, the conformational change of the α3 helix and β3-β4 hairpin in site V is vital to expose the fusion peptide and initiate the fusion process. Antibodies targeting site V likely lock the fusion peptide and prevent the formation of the long helical bundles that are present in the post-fusion conformation. However, based on current findings, the frequency of hMPV site V-specific antibodies is relatively low compared to antibodies targeting site III, site IV, and DS7 site. Therefore, it would be important to consider boosting antigenic site V targeting antibodies in hMPV F-based vaccines.

## Materials and Methods

### Ethics statement

This study was approved by the University of Georgia Institutional Review Board. Healthy human donors were recruited to the University of Georgia Clinical and Translational Research Unit and written informed consent was obtained. Animal studies were approved by the University of Georgia IACUC.

### Blood draws and PBMC isolation

After obtaining informed consent, 90 mL of blood was drawn by venipuncture into 9 heparin-coated tubes, and 10 mL of blood was collected into a serum separator tube. Peripheral blood mononuclear cells (PBMCs) were isolated from human donor blood samples using Ficoll-Histopaque density gradient centrifugation, and PBMCs were frozen in the liquid nitrogen vapor phase until further use.

### Production and synthesis of recombinant hMPV F protein

hMPV A1, A2, B1, B2 F and hMPV B2F-GCN4 recombinant proteins were synthesized from the plasmids obtained from GenScript cloned into pcDNA3.1+ vector. They were expanded by transforming them into DH5α cells against ampicillin (Thermo Scientific) resistance 100 ug/ml. The plasmids were purified using E.N.Z.A. plasmid maxiprep kit (Omega BioTek) following the manufacturer’s instructions. 1 mg of plasmid was mixed with 4 mg of polyethyleneimine (PEI; PolySciences Inc.) in Opti-MEM cell culture medium (Thermo Scientific) and incubated for 30 minutes. This was followed by addition of the DNA-PEI mixture to 10^6^ cells/ml 293 cells in Freestyle 293 expression medium (Thermo Fischer). After 5 days of incubation, the cultures were centrifuged at 6000 g to pellet the cells. The supernatant was filtered through a 0.45 μm sterile filter. The recombinant proteins were purified directly by affinity chromatography, HisTrap Excel columns (GE Healthcare Life Sciences). Prior to loading the supernatant onto the column, it was washed with 5 column volumes (CV) of a wash buffer containing 20 mM Tris-HCl (pH 7.5), 500 mM NaCl and 20 mM Imidazole. After passing the supernatant through the column, it was washed with the same wash buffer (5 CV) to reduce non-specific binding and finally eluted with buffer containing 20 mM Tris-HCl pH 7.5, 500 mM NaCl and 250 mM Imidazole. After elution, the proteins were concentrated with Amicon Ultra-15 centrifugal units with a molecular cut-off of 30 KDa (Sigma).

### Trypsinization of hMPV F

In order to obtain trimeric hMPV F, TPCK (L-1-tosylamido-2-phenylethyl chloromethyl ketone)-trypsin (Thermo Scientific) was dissolved in double-distilled water (ddH_2_O) at 2 mg/mL. hMPV B2 F obtained, was incubated with 5 TAME (p-toluene-sulfonyl-L-arginine methyl ester) units/mg of TPCK-trypsin for 1 hr at 37°C. The trimeric and monomeric fractions of hMPV F were separate by size exclusion chromatography on a Superdex S200, 16/600 column (GE Healthcare Life Sciences) in column buffer (50 mM Tris pH 7.5, and 100 mM NaCl). Both fractions were separated by their unique elution profiles. Once separated, they were concentrated as mentioned earlier. Post-fusion hMPV F was obtained by heating the pooled trimeric fractions at 55 °C for 20 minutes on a water bath to induce post-fusion conformation.^29^

### Generation of hMPV F-specific hybridomas

For hybridoma generation, 10 million peripheral blood mononuclear cells purified from the blood of human donors were mixed with 8 million previously frozen and gamma irradiated NIH 3T3 cells modified to express human CD40L, human interleukin-21 (IL-21), and human BAFF^46^ in 80 mL StemCell medium A (StemCell Technologies) containing 6.3 μg/mL of CpG (phosphorothioate-modified oligodeoxynucleotide ZOEZOEZZZZZOEEZOEZZZT; Invitrogen) and 1 μg/mL of cyclosporine (Sigma). The mixture of cells was plated in four 96-well plates at 200 μl per well in StemCell medium A. After 6 days, culture supernatants were screened by ELISA for binding to recombinant hMPV B2 F protein, and cells from positive wells were electrofused as previously described.^46^ Cells from each cuvette were resuspended in 20 mL StemCell medium A containing 1× HAT (hypoxanthine-aminopterin-thymidine; Sigma-Aldrich), 0.2× HT (hypoxanthine-thymidine; Corning), and 0.3 μg/mL ouabain (Thermo Fisher Scientific) and plated at 50 μl per well in a 384-well plate. After 7 days, cells were fed with 25 μl of StemCell medium A. The supernatant of hybridomas were screened after 2 weeks for antibody production by ELISA, and cells from wells with reactive supernatants were expanded to 48-well plates for 1 week in 0.5 mL of StemCell medium E (StemCell Technologies), before being screened again by ELISA. Positive hybridomas were then subjected to single-cell fluorescence-activated sorting into 384-well plates containing 75% of StemCell medium A plus 25% of StemCell medium E. Two weeks after cell sorting, hybridomas were screened by ELISA before further expansion of wells containing hMPV F-specific hybridomas.

### RT-PCR for hybridoma mAb variable gamma chain and variable light chain

RNA was isolated from expanded hybridoma cells using the ENZA total RNA kit (Omega BioTek) according to the manufacturer’s protocol. A High-Capacity cDNA Reverse Transcription Kit (Applied Biosystems) was used for cDNA synthesis. Three separate sets of primer mixes were used for nested PCR to amplify the variable regions of gamma, kappa, and lambda chains.^48^ The products from the second PCR were analyzed by agarose gel electrophoresis and purified PCR products (ENZA cycle pure kit; Omega Biotek) were submitted to Genewiz for sequencing. Sequences were analyzed using IMGT/V-Quest.^49^

### Antigen-specific single B cell sorting and expression of recombinant mAbs

Ten million human PBMCs were washed twice with FACS buffer and then resuspended with 1 mL FACS buffer. The cells were treated with 5% Fc receptor blocker (BioLegend) for 30 minutes and then stained with following antibodies: human CD19-APC, human IgM-FITC, human IgD-FITC, Ghost Dye™ Red 710, and PE/BV605-streptavidin conjugated hMPV B2 F. Antigen specific B cells were gated with CD19+/IgM-/IgD-/Ghost dye-/PE+/BV605+ and sorted in catch buffer B (Qiagen TCL Buffer + 1% beta mercaptoethanol) by one cell per well in a 96 well plate. Sorted cells were flash frozen and stored in −80 °C until they were used for RNA extraction. The RNA was extracted with Agencort RNAClean XP kit, SPRI Beads (Beckman Coulter) and immediately reverse transcribed to cDNA with SuperScript IV Synthesis System (ThermoFisher). The variable region sequences of IgG heavy/light chains were determined by nested PCR as described above. Based on the usage of V/D/J gene alleles, cloning PCR primers were picked for cloning PCR with the first PCR products as the template. Purified cloning PCR products of heavy/light chains were cloned into expression vectors (AbVec-hIgG1, AbVec-hIgKappa, and pBR322-based Ig-lambda expression vector) and the plasmids were sent to Genewiz for sequencing. After confirming all the sequences are correct, the HC/LC plasmids were transformed into DH5α for plasmid maxiprep. Recombinant mAbs were expressed by transfecting 293 cells with HC/LC plasmids and purified from culture supernatant with Protein G column (Cytiva).

### Enzyme-linked immunosorbent assay for binding to hMPV F protein

The 384 well plates (catalog number 781162; Greiner Bio-One) used for ELISA were coated with 2 μg/ml (in PBS) of the recombinant protein (antigen) and incubated overnight at 4°C. This was accompanied by washing the plates once, with water followed by blocking them for 1 hr at room temperature with Block buffer comprising of 2% milk supplemented with 2% goat serum in PBS and 0.05% Tween 20 (PBS-T). The plates were washed again thrice with PBS-T. 25 μl of the serially diluted primary antibodies were added to the wells and incubated for an hour at room temperature followed by three washes with PBS-T. Goat anti-human IgG Fc secondary antibody (Southern Biotech) diluted in block buffer (1:4000) was next applied to the wells and incubated again at room temperature for an hour. The plates were washed again with PBS-T thrice and 25 μl of PNPP (p-nitrophenyl phosphate) diluted to a concentration of 1 mg/ml in a buffer containing 1M Tris base and 0.5 mM Magnesium Chloride having a pH of 9.8 was added. Prior to reading the absorbance at 405 nm on a Bio Tek plate reader, the plates were incubated one last time at room temperature for 1 hr. The binding assay data was analyzed on GraphPad Prism using a nonlinear regression curve fit and log(agonist)-versus-response function for calculating the EC_50_ values.

### hMPV plaque neutralization experiments

LLC-MK2 cells used for this experiment were grown in Opti-MEM I (Thermo Fischer Scientific) that was supplemented with 2% fetal bovine serum in T225 cell culture flasks (catalog number 82050-870) at 37 °C in a CO_2_ incubator. Two days before beginning the neutralization assay, 40,000 cells/well were plated on 24-well plates. Serially diluted sterile-filtered mAbs isolated from hybridoma supernatants were added to the suspension of either of hMPV strains, CAN/97-83 and TN/93-32 in equal amounts (1:1) and incubated for 1 hour on the day of the experiment. This was followed by addition of 50 μl of the virus-antibody mixture to the LLC-MK2 cells after washing of the excess FBS from the OPTI-MEM media with PBS three times. The mixture was incubated at room temperature for an hour with constant rocking. The cells were next overlaid with 0.75% methylcellulose dissolved in Opti-MEM I supplemented with 5 μg/ml of trypsin-EDTA and 100 ug/ml of CaCl_2_. The cells were incubated for 4 days and fixed with 10% neutral buffered formalin. Cell monolayers were next blocked with block buffer comprising of 2% nonfat milk supplemented with 2% goat serum in PBS-T for an hour. Next, the plates were washed thrice with water and 200 μl of MPV 364 was added to a final concentration of 1 μg/ml (1:1000 dilution) in blocking solution. The plates were then washed three times with water and 200 μl of goat anti-human IgG HRP secondary antibody (Southern Biotech) diluted to a ratio of 1:2000 in block buffer was added and incubated for 1 hr at room temperature followed by an hour of incubation. Plates were washed again with water five times and 200 μl of TrueBlue peroxidase substrate (SeraCare) was added to each well. The plates were incubated for 20-30 minutes until the plaques were clearly visible. Plaques were counted manually under a microscope and compared to the virus-only control. GraphPad Prism was used to calculate the IC_50_ values using a nonlinear regression curve fit and the log(inhibitor)-versus-response function.

### Epitope Binning

100 μg/ml of the his-tagged hMPV B2F (not trypsin treated) protein was immobilized on anti-penta-His biosensor tips (Forte’ Biosciences) for 120 s after obtaining the initial baseline in running buffer (PBS, 0.5% BSA, 0.05% Tween 20 and 0.04% thimerosal). Base line was measured again with the tips immersed in wells containing 100 μg/ml of the primary antibody for 300 s. This was followed by immersing the biosensor tips again for 300 s in the secondary antibody at 100 μg/ml. Binding of the second mAb in the presence of the first mAb as determined by comparing the maximal signal of the second mAb after the first mAb was added to the maximum signal of the second mAb alone. Non-competing mAbs were those whose binding was greater than or equal to 70 % of the uncompeted binding. Between 30% and 60% was considered intermediate binding and anything lower that 30% was considering as competing for the same site.

### Antibody-dependent phagocytic activity of mAbs

To measure antibody-dependent phagocytic activity, 2×10^9^ 1-μm Neutravidin-coated yellow-green FluoSpheres (Invitrogen #F8776) were resuspended in 1 mL of 0.1% PBS. The FluoSpheres were then centrifuged at 5000 rpm for 15 minutes, 900 μL supernatant was removed, and the FluoSpheres were resuspended with 900 μL of 0.1% PBS. This process was repeated for a second wash, then the FluoSpheres were resuspended with 20 μg of biotinylated hMPV B2 F protein. The FluoSpheres were then incubated overnight at 4 °C, protected from light, with end-to-end rocking. Next, hMPV F-specific antibodies were diluted in complete RPMI media (cRPMI, RPMI + 10% FBS) to a final concentration of 1 μg/mL in a U-bottom 96-well plate. Then, 20 μL of antibody dilution was transferred into a clean F-bottom 96-well plate, and 10 μL of FluoSpheres were added with the antibody followed by a 2 hr incubation at 37 °C for opsonization. After 1.5 hr, THP-1 cells were centrifuged at 200 × g for 5 min, washed once with PBS, then resuspended in culture medium (RPMI & 10% FBS) at a concentration of 5×10^5^ cells/mL. Then, 200 μL of cells were added to each well and incubated at 37 °C with 5% CO_2_ while shaking for 6 hr. Once the incubation finished, the plate was then centrifuged at 2000 rpm for 5 min. Then, 100 μL was pipetted out of each well and replaced with 100 μL of cold 4% paraformaldehyde to fix the cells. The plate was then left at room temperature for 20 min, protected from light. The plate was then stored at 4 °C in the dark. Cells were then analyzed with a NovoCyte Quanteon flow cytometer. The percentage of fluorescent beads containing THP-1 cells in each sample (% phagocytosis) was used to calculate % increase vs. no mAb control. The phagocytic scores were calculated as previously described^50^ (geometric mean intensity - the geometric mean intensity of the no mAb control) × % phagocytosis.

### Animal studies

BALB/c mice (6 to 8 weeks old; The Jackson Laboratory) were randomly selected to each group that contains 5 males and 5 females. All the mice were pre-bleed before the study to verify the mice were not pre-exposed to hMPV by ELISA. Each mouse was intranasally infected with hMPV TN/93-32 (5×10^5^ PFU) and euthanized five days post-infection. Mice were i.p. injected with PBS/MPV467/control antibodies (10 mg/kg) 24 hours prior to infection (prophylaxis) or three days post-infection (treatment). At the end point, serum was collected for ELISA to determine the presence of mAb MPV467, the lungs were collected and homogenized to determine the viral load through immunostaining as described above.

### Recombinant protein production for cryo-EM studies

The prefusion hMPV F construct DS-CavEs2-IPDS (hMPV F A1 NL/1/00, residues 1-490) used for structural studies includes the previously described G294E, A185P, L219K, V231I, E453Q and furin cleavage site substitutions^31^ (and Hsieh et al, *in press*). Disulfide substitutions included are L110C/N322C, T127C/N153C, A140C/A147C and T365C/V463C (Hsieh et al, *in press*) as well as an interprotomer disulfide at V84C/A249C.^51^ DS-CavEs2-IPDS was cloned into the mammalian expression vector pαH with a C-terminal “GGGS” linker sequence followed by the T4 fibritin trimerization motif “foldon”^52,53^ an HRV3C protease site, an 8xHis tag, and a Strep-TagII.^31^ Transient co-transfection of FreeStyle 293F cells (ThermoFisher) at a 4:1 ratio of DS-CavEs2-IPDS:furin-expressing plasmids by polyethylenimine (PEI) was used for protein expression. Kifunensine and Pluronic F-68 (Gibco) were introduced three hours post-transfection to a final concentration of 5 μM and 0.1% (v/v), respectively. Six days post-transfection, Strep-Tactin Sepharose resin (IBA) was used to purify soluble protein from the cell supernatant which had been filtered and buffer-exchanged into PBS by tangential flow filtration. Buffer containing 100 mM Tris pH 8.0, 150 mM NaCl, 1 mM EDTA and 2.5 mM desthiobiotin was used to elute the strep-tagged protein. After concentrating the protein using a 30 kDa molecular weight cut-off Amicon Ultra-15 centrifugal filter unit (Millipore) the protein was further purified by size-exclusion chromatography using a Superose 6 Increase 10/300 column (GE Healthcare) in 2 mM Tris pH 8.0, 200 mM NaCl, and 0.02% NaN_3_ running buffer.

### Cryo-EM Sample Preparation and Data Collection

Purified DS-CavEs2-IPDS was combined with a 1.5-fold molar excess of MPV467 Fab incubated at room temperature for 10 minutes before being moved to ice. Just before freezing, sample was diluted to a concentration of 0.66 mg/mL hMPV F in 2 mM Tris pH 8.0, 200 mM NaCl, and 0.02% NaN_3_ buffer. Then 1 μL of 0.5% amphipol A8-35 was combined with 10 μL of diluted sample and 4 μL of this sample was applied to a gold 1.2/1.3 300 mesh grid (Protochips Au-Flat) that had been plasma cleaned for 180 seconds using a Solarus 950 plasma cleaner (Gatan) with a 4:1 ratio of O_2_/H_2_. Grids were plunge-frozen using a Vitrobot Mark IV (Thermo Fisher) with a 10 °C, 100% humidity chamber. Blotting settings were 5 seconds of wait followed by 4 seconds of blotting with −2 force before plunging into nitrogen-cooled liquid ethane. Using a Glacios (Thermo Scientific) equipped with a Falcon 4 direct electron detector (Thermo Scientific), a single grid was imaged to collect a total of 1,458 images. Data were collected at a 30° tilt with magnification of 150,000x corresponding to a calibrated pixel size of 0.94 Å/pix and a total exposure of 40 e^−^/ Å^2^. Data collection statistics are listed in **Table S2**.

### Cryo-EM data processing

Micrographs were corrected for gain reference and imported into cryoSPARC Live v3.2.0 for initial data processing: motion correction, defocus estimation, micrograph curation, particle picking and extraction, and particle curation through iterative streaming 2D class averaging. 2D averages were used to generate templates and template-based particle picking was carried out. Curated particles were exported to cryoSPARC v3.2 for further processing via rounds of 2D classification, ab initio reconstruction, heterogeneous refinement, homogenous refinement, and non-uniform homogenous refinement using C3 symmetry. Masking and particle subtraction were used for further non-uniform refinement. Finally, the particle-subtracted non-uniform refinement map was sharpened using DeepEMhancer.^45,54^ EM processing workflows are shown in **Figure S3**, and EM validation results are shown in **Figure S4**. For model building, an initial hMPV F model was generated from PDB ID: 5WB0 and the crystal structure of MPV467 Fab which were used to dock into the cryoEM maps using UCSF ChimeraX.^55^ Models were built further and iteratively refined using a combination of Coot^56^ PHENIX^57^ and ISOLDE.^58^ Model statistics are shown in **Table S2**.

## Acknowledgements

These studies were supported by National Institutes of Health grants 1R01AI143865 (JJM) and 1K01OD026569 (JJM). This work was funded in part by Welch Foundation grant number F-0003-19620604 (JSM), and the Georgia Research Alliance (RAT). We thank the University of Georgia Clinical and Translational Research Unit for assistance with human subject identification and blood draws, and the University of Georgia Center for Tropical and Emerging Global Diseases flow cytometry core for assistance with cell sorting. We acknowledge the University of Texas College of Natural Sciences and award RR160023 of the Cancer Prevention and Research Institute of Texas for support of the EM facility at the University of Texas at Austin. F.R was supported by national Institutes of Health NIGMS grant GM109435, Post-Baccalaureate training in Infectious diseases Research.

## Data deposition

The map and coordinates for the structure of MPV467 in complex with the hMPV F protein were deposited in the protein data bank under accession ID ####.

