## Supplemental Figures and Tables for "Structural basis for ultrapotent neutralization of human metapneumovirus"

| **Table S1. Monoclonal antibody sequences** | | | |
|---|---|---|---|
| mAb | Subject | HC sequence AA | LC sequence AA |
| MPV 467 | 3 | QVQLVESGGGVVRPGTSLRVSCAAFDFNFRDYGMHWVRQAPGKGLEWVAGIWYDGSNKDYADSVKGRFTISRDNSQNTLYLQMNSLRVEDTAVYYCARDPRTHREGALSHFDSWGQGTLVTVSS | PNELTQDPAVSVALGQTVRITCQGDSLRNYFAGWYQQKPGQAPLLVLYGENIRPSGIPDRFSGSSSGNTVSLTITGAQAEDEADYYCNSRDNSGNHWVFGGGTRLTVL |
| MPV 487 | 8 | QVQLLESGGGLVQPGRSLRLSCTASGFTFDDYAMHWVRQLPGKGLEWVSGISWNSDNIGYADSVKGRFTISRDNGKNSLYLQMNSLRAEDTAFYYCAKDVRIDYDLLIGHIDYWGPGTLVTVSS | TQSPVTLSLSPGERATLSCRASQSFGGYCAWYQQKPGQSPRLLIYDASNRATGVPARFSGSGSGTDFTLTISSLEPEDSAVYFCQQRGNWPIFGQGTRLEIK |
| MPV 454 | 2 | QVQLVESGGGLVKPGGSLRLSCSTSGFVFSSYAMHWVRQAPGKGLEHVSAISSTGDNTHYADSVKGRFTISRDNSRGRLYLQMTSVRPEDAALYYCVKDLYSGRFYYFLDDWGQGTRIIVSS | DIQVTQSPSSLSASVGDRVTISCRASQNISNYLNWFQQRPGKAPKLLIYVASTLQSGAPSRFSGSGSGTDFTLTITDLQPEDFATYYCQQSYNTPPITFGQGTRLEIK |
| MPV 482 | 6 | QVQLVQSGGGVVQPGRSLRLSCAASGFTFSSFAMHWVRQAPGKGLEWVALISDDGNNKYHADSVKGRFTISRDSSKNTLYLQMNSLRGEDTAVYYCARDGSVVGAAAHCDHWGQGTLVTVSS | SYELTQPPSVSVSPGQTARITCSGDALPNQCAYWYQQKPGLAPVLLIYKDKERPSGIPERFSGSISGTTVTLTISGVQAEDEADYYCQSSDSSSTVVFGGGTKLTVL |
| MPV 486 | 7 | QVQLVESGGHLVQSGGSLRLSCAASGFTFSNYILNWVRQAPGKGLEWISYITGGSSAIYYADSVKGRFTISRDDAKNSLYLQMNNLRAEDTAVYYCARSHGYDSSGYYYYFAMDVWGQGTTVTVSS | QSVLTQPPSASGTPGQRVTISCSGSNSNIESNTVNWYQQIPGTAPKLLIYSDNRRPSGVPDRFSGSKSGTSASLAISGLQSEDEADYYCAAWDDSLIGYVFGTGTKVTV |
| MPV 478 | 4 | QVQLVQSGAEVKKPGASVKVSCNASGYTFTGYYIHWVRQAPGQGLEWMGWINPRSGGTNYAQKFQGRVTLTRDTSITTAYMDVTRLRPDDTAVYYCATTREGIVLMPRGRGDDALDTWGQGTTVTVSS | DIQMTQSPVSLTVSLGERATINCKSSQSVLSTSNNENYLAWYQQKPGQPPNLLIYWASTRESGVPDRFSGSGSGTDFTLTISSLQAEDVAVYYCQQYFATPITFGPGTKVDVK |
| MPV 483 | 6 | QVQLVQSGAELKKPGSSVKVSCKASGDTFSNYAISWVRQAPGQGLEWMGGIIPIYNTANYAQKFQGRVTLTADESTSTAYMELSSLRSEDTAVYYCARDVRNNWSVLRGARYYYYGMDVWGQGTTVTVSS | IRLMTQTPFSLSASVGDRVTVTCRASQDISNSLAWFQQKPGKAPKSLIYAASSLQSGVPTRFSGGGSGTDFTLTISSLQPEDFATFYCLQYDSFPPTFGGGTKVEIK |
| MPV503 | 5/6 | QVQLVQSGAEVKKPGSSVKVSCKASGGTFSSYTFSWVRQAPGQGLEWMGGIIPIFGTANYAQKFQDRVTITADESTSTAYMELSSLRSEDTAVYYCARGRVVVIASHYYYYDMDVWGQGTTVTVSS | VSLSVSPGERATLSCRASQSVSSNLAWYQQKPGQAPRLLIYGASTRATGIPARFSGSGSGTEFTLTISSLQSEDFAVYYCQQYDNWPPLTFGGGTKVEIK |
| MPV 481 | 5 | QVQLVQSGGGLVKPGGSLRLSCAASGLPFNNAWMNWVRQAPGKGLEWVGRIKSKIDGGTTEYAAPVKGRFTISRDDSENTLYLQMNSLRTEDTAVYYCTTEYYALLSGNYSDCWGQGTLVTVSS | IRLMTQSPFSLSLSPGERATLSCRASQSIKRAYLGWYQQKPGQAPRLLIYGASNRATGIPDRFSGSGSGTDFTLTISRLEPEDFAVYYCQQYGDSPGGSFGQGTKLEIR |
| MPV 464 | 2 | QVQLQESGPGLVKPSQTLSLTCTVSGGSVRGGGDYWSWIRRPPGKGLEWIGHIYNSGTTFYNPSLKSRVTISIDTSKNQFSLKLRSVTAADTAVYYCGRVGSRATGLPDWIDPWGQGTLVTVSS | SYELTQPPSVSVSPGQTATITCSGTNLGAKYSCWYQQKPGQSPVLVIFQDNKRASGIPARFSASTSGDTATLTISGTQVMDEADYFCQAWDSRTAVFGGGTTLTV |
| MPV414 | 1 | QVQLVQSGPEVKKPGSSVMVSCKTSGGNFNNFAISWVRQAPGQGLEWMGGIVPIFGTANYAQKSQDRVTITADPSSSTAY | DIVMTQSPATLSLSPGERATLSCRASQSVGDKLAWYQQKPGQAPRLLIYGASTRATDIPARFSGSGSGTEFTLT |

| | | | |
|---|---|---|---|
| | | LQLSSLTSDDTAIYYCARDLGRSSNYYY YSVFESWGQGTLVTVSS | VSSLQPEDFAVYYCQQYENWPPIT FGQGTRLEIK |
| MPV 456 | 2 | QVQLQESGPGLVKPSQTLSLTCTVSGG SVRGGGDYWSWIRRPPGKGLEWIGHIY NSGTTFYNPSLKSRVTISIDTSKNQFSLK LRSVTAADTAVYYCGRVGSRATGLPDW IDP | TLSASVGDRVTITCRASQSISSWLA WFQQKPGKAPKLLIYKTSSLESGV PSRFSGSGSGTEFTLTISSLQPDDF ATYYCQEYNSYSRTFGQGTKVEIK |
| MPV86 | 1 | QVQLVQSGAEVKKPGSSVKVSCKASGD TFNDVALSWVRQAPGQGLEWLGGIIPF FGTANYAQRFQDRVTITADASTSTAYLE LGGLRSEDTAVYYCARGWSCRNITCFM MGYYFYIMDVWGQGTLVTVSS | LSLSPGERATLSCRASQSVGNFLA WYQQKPGQTPRLLIQDASNRATGI PARFSGTGSGTDFTLTISSLEPEDF AVYYCQQRSNWPPTFGPGTKVDIK |
| MPV 491 | 9 | QVQLQESGPGLVKSSQTLSLTCTVSGG SISSVTSHWTWIRQHPGKGLEWIGYIFS SGTTYYSPSLRSRLTMSVDTSKNQFSL QLSSVTAADTAMYYCARGIYCGDSCYK GTDYWGRGTLVTVSS | IRLMTQTPVTLPVTPGEPASISCRS SQSLLHSNGYNYLDWYLQKPGQS PQLLIYRGSTRASGVPDRFSGSGS GTEFTLKISRVEAEDVGVYYCMQG LQTPYTFGQGTKLESR |
| MPV 489 | 8 | QVQLLESGGGLVQPGGSLRLSCAASKF TFNNYEMNWLRQAPGKGLEWVSSISSS GDTIHNADSVKGRFIISRDNAKNSLHLQ MNGLRAEDTAVYYCARAGSGWAYDAF DIWGLGTMVTVSS | LSASVGDRVTITCRASQSISSFLNW YQQKPGKAPKLLMYAASSLPVGVP SRFSGSGSGPEYTLTISNLQPEDFA TYYCQQGYSSPPTFGGGTRLEVK |
| MPV 488 | 8 | QVQLVQSGFEVKKPGASVKVSCKASGY NFNNYGISWVRQAPGQGLEWMGWISA YTGNTNEAQKFQGRVSMTTDTSTSTAY MEVRSLRSDDTAVYYCARDIGSSMFYS YFYGMDVWGQGTTVIVSS | SLSVSLGERATINCKSSESVLYNSN NENYLDWYQQKPGQPPKLLIYWAS TRASGVPDRFSGSASGTDFTLTISS LQAEDVAVYYCQQYYSTPRTFGQ GTKVEIK |
| MPV 485 | 3 | QVQLQEPGPGLVKPSETLSLTCTVSGG SISSTNSFWGWVRQPPGKGLEWIGSIY YSGTTYYNSSLKSRVTISVDTSKNQFSL RLSSVTAADTAVYYCARQGLTSSWYDG SGLDVWGRGTKVTVSS | SYELTQPPSVSVSPGQTASITCSG DKLGNKYACWYQQRPGQSPVLVIY QDTKRPSGIPERFSGSNSGNTATL TISGTQAMDEADYYCQAWDSNTV VFGGGTKLTVL |
| MPV 477 | 4 | QVQLVQSGAEVKKPGASVEVSCKTSGY TFTXYYLHWVRQAPGQGLEWMGWINP RSGGTKYGQKFQGRVTMTRDTSISTAY MDLRGLRSDDTAVYYCARAPLLTVYAV AHRSGENRFDPWGQGTVV | QPPSLSGAPGQTVTISCTGSGSNI GAGYDVNWYQCLPGTAPKLLMFD NSNRPSGVPDRFSGSRSGASASL AITGLQAEDEADYYCQSYDSGLSG WVFGGGTKLTVL |

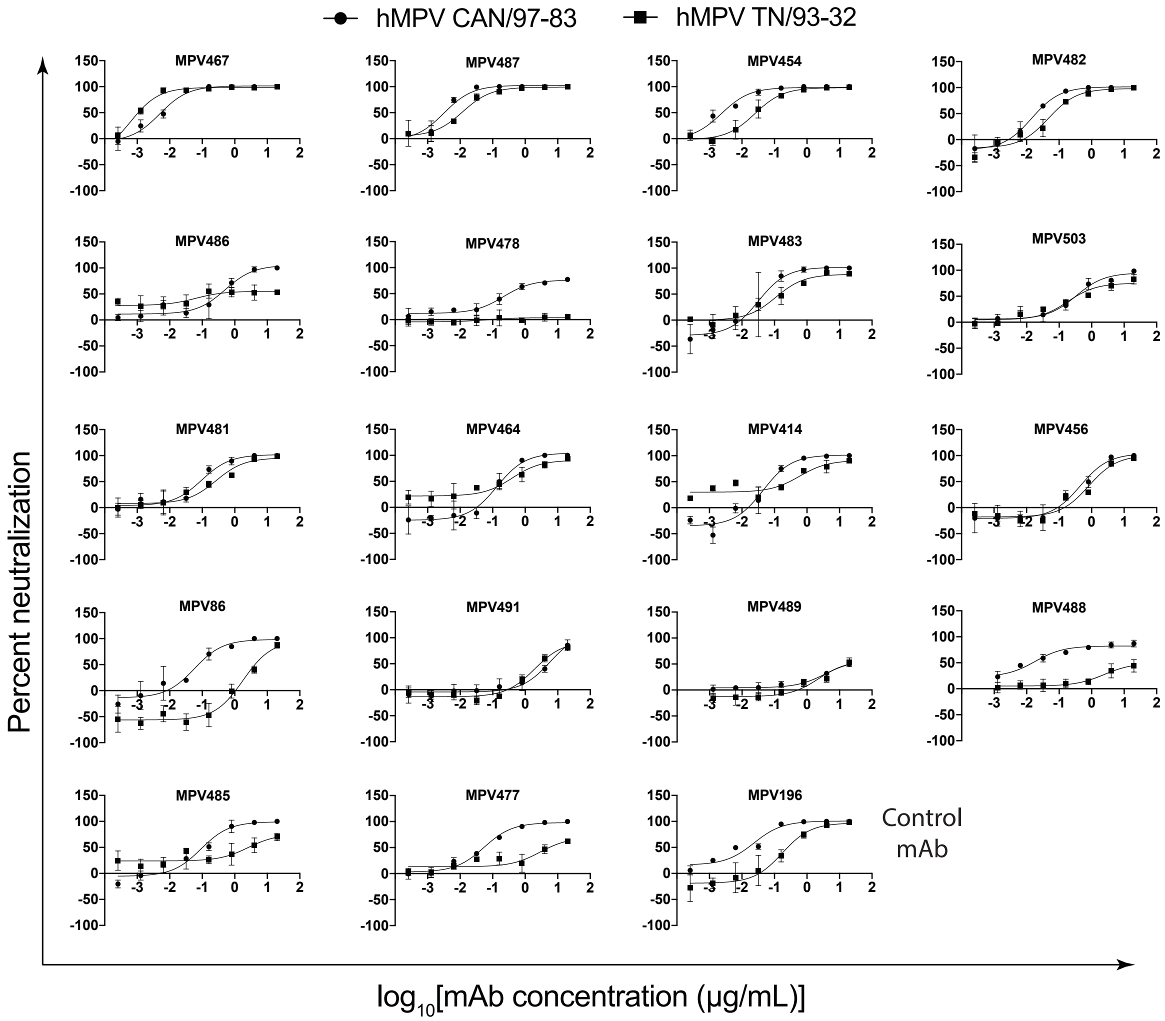

**Figure S1. Neutralization profiles of the hMPV F protein-specific mAbs.** Data represent the averages from three replicates, and error bars are the standard deviation. Data are representative of results from at least two independent experiments.

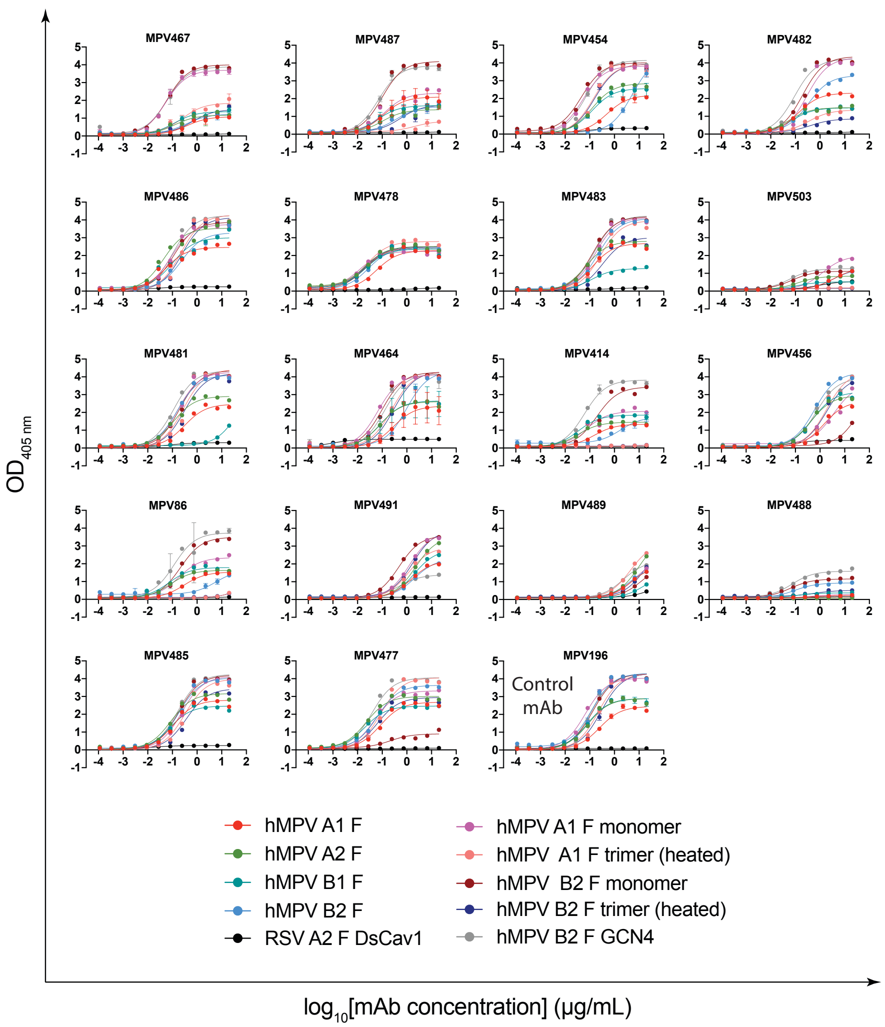

**Figure S2. ELISA binding curves of the hMPV F protein-specific mAbs.** ELISA binding curves of the isolated mAbs with recombinant hMPV F protein constructs. Prefusion RSV F protein (DsCav1) was utilized to determine if any mAb cross-reacts with RSV F. The previously discovered hMPV F-specific mAb MPV196 was used as a negative binding control. Each point represents the average of data from four replicates, and error bars represent the standard deviation. Data are representative of results from at least two independent experiments.

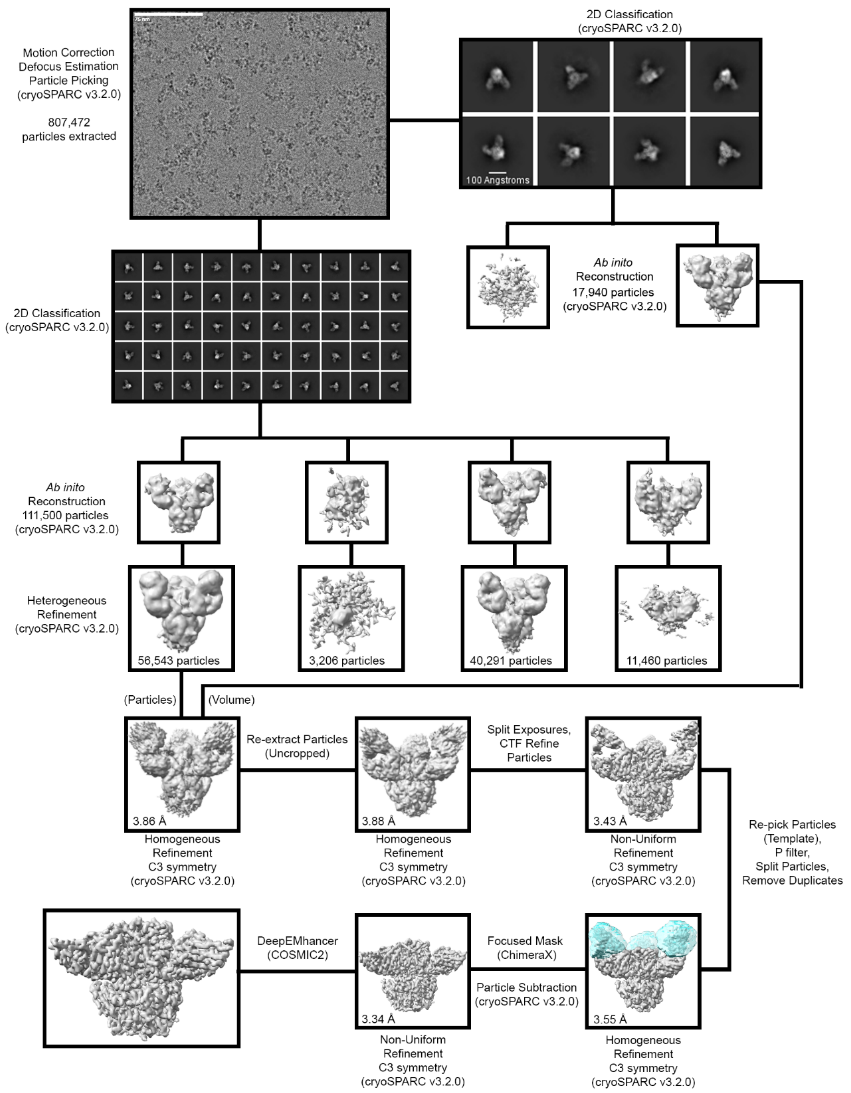

**Figure S3. MPV467 cryo-EM processing workflow.** Each step, from representative micrograph to DeepEMhancer map, of the cryo-EM data processing workflow is shown. Computational programs and algorithms used are labeled for each step. The mask used for particle subtraction is colored as a transparent cyan.

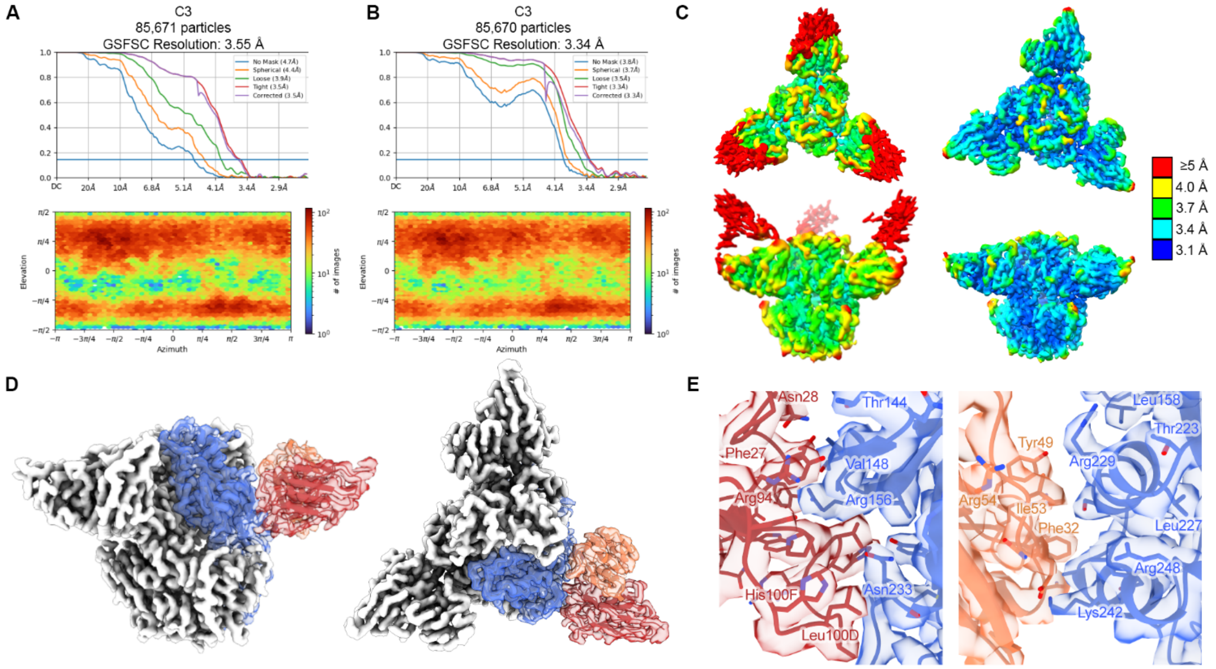

**Figure S4. MPV467 cryo-EM Validation.** (A) (top) FSC curves for the homogeneous refinement 3D reconstruction. Horizontal blue line corresponds to an FSC value of 0.143. (bottom) Viewing distribution plot calculated in cryoSPARC. (B) FSC curves and viewing distribution plot for the particle-subtracted non-uniform refinement 3D reconstruction. (top) The FSC curves for the non-uniform refinement 3D reconstruction. The horizontal blue line corresponds to an FSC value of 0.143. (bottom) Viewing distribution plot calculated in cryoSPARC. (C) Cryo-EM maps of homogenous refinement (left) and particle-subtracted non-uniform refinement (right) colored by local resolution. Cryo-EM maps are shown as top (top) and side (bottom) views. (D) The cryo-EM map of DS-CavEs2-IPDS bound by MPV467 is shown in a side view (left) and top view (right). A single protomer is shown as a transparent colored surface with a docked ribbon model (DS-CavEs2-IPDS: blue; MPV467 heavy chain: red, MPV467 light chain: orange). (E) The binding interface for MPV467 heavy chain (left) and light chain (right) with DS-CavEs2-IPDS. Cryo-EM map is shown as a transparent surface with the docked model shown as ribbon and sticks. Colored the same as in D.

### Supplemental Table 2. EM data collection for MPV467

**EM data collection**

| | |
|---|---|
| Microscope | Glacios |
| Voltage (kV) | 200 |
| Detector | Falcon 4 |
| Magnification (nominal) | 150,000 |
| Pixel size (Å/pix) | 0.94 |
| Exposure rate ($e^-$/pix/sec) | 2.55 |
| Exposure ($e^-/Å^2$) | 40 |
| Defocus range (mm) | 1.0-2.5 |
| Tilt angle (˚) | 30 |
| Micrographs collected | 1,458 |
| Micrographs used | 1,114 |
| Particles extracted (total) | 807,472 |
| Automation software | SerialEM |
| Sample | hMPV F + MPV467 Fab |

**3D reconstruction statistics**

| | Overall | hMPV F + MPV467 Fab |
|---|---|---|
| Particles | 85,671 | 85,670 |
| Symmetry | C3 | C3 |
| Map sharpening B-factor | -122.3 | -113.3 |
| Unmasked resolution at 0.5 FSC (Å) | 7.7 | 4.2 |
| Masked resolution at 0.5 FSC (Å) | 4.0 | 3.7 |
| Unmasked resolution at 0.143 FSC (Å) | 4.7 | 3.8 |
| Masked resolution at 0.143 FSC (Å) | 3.55 | 3.34 |

**Model refinement and validation statistics**

| | |
|---|---|
| Refinement package | PHENIX |
| Refinement tool | Real-space refinement |

| | |
|---|---|
| Refinement strategies | min global, local_grid_search, adp, ss restraints, rotamer restraints, Ramachandran restraints |
| Composition | |
| Amino acids | 1,973 |
| RMSD bonds (Å) | 0.006 |
| RMSD angles (°) | 0.60 |
| Average B-factors | |
| Amino acids | 80.3 |
| Ramachandran | |
| Favored (%) | 96.4 |
| Allowed (%) | 3.6 |
| Outliers (%) | 0.0 |
| Rotamer outliers (%) | 0.72 |
| Clash score | 3.77 |
| C-beta outliers (%) | 0 |
| CaBLAM outliers (%) | 1.77 |
| 0.5 FSC model (Å) | 3.5 |
| CC (mask) | 0.85 |
| MolProbity score | 1.40 |
| EMRinger score | 3.46 |
